# Neural adaptation to the eye’s optics through phase compensation

**DOI:** 10.1101/2024.08.21.608968

**Authors:** Antoine Barbot, John T. Pirog, Cherlyn J. Ng, Geunyoung Yoon

**Affiliations:** Flaum Eye Institute, University of Rochester Medical Center, Rochester NY, United States; Center for Visual Science, University of Rochester, Rochester NY, United States; Institute of Optics, University of Rochester, Rochester NY, United States; College of Optometry, University of Houston, Houston TX, United States

**Keywords:** phase perception, adaptation, optical blur, neural compensation, adaptive optics, keratoconus

## Abstract

How does the brain achieve a seemingly veridical and ‘in-focus’ perception of the world, knowing how severely corrupted visual information is by the eye’s optics? Optical blur degrades retinal image quality by reducing the contrast and disrupting the phase of transmitted signals. Neural adaptation can attenuate the impact of blur on image contrast, yet vision rather relies on perceptually-relevant information contained within the phase structure of natural images. Here we show that neural adaptation can compensate for the impact of optical aberrations on phase congruency. We used adaptive optics to fully control optical factors and test the impact of specific optical aberrations on the perceived phase of compound gratings. We assessed blur-induced changes in perceived phase over three distinct exposure spans. Under brief blur exposure, perceived phase shifts matched optical theory predictions. During short-term (∼1h) exposure, we found a reduction in blur-induced phase shifts over time, followed by after-effects in the opposite direction–a hallmark of adaptation. Finally, patients with chronic exposure to poor optical quality showed altered phase perception when tested under fully-corrected optical quality, suggesting long-term neural compensatory adjustments to phase spectra. These findings reveal that neural adaptation to optical aberrations compensates for alterations in phase congruency, helping restore perceptual quality over time.

---

How we perceive the world is intrinsically constrained by our sensory receptors. The optics of the human eye degrade the quality of the images projected onto the retina, placing fundamental limits on visual processing (Artal et al., 2004; Webster, 2015; Webster et al., 2002). Optical blur affects the information transmitted to the retina in two ways: it attenuates the contrast of the retinal image and disrupts the spatial phase (position) of visual signals relative to each other (**Fig.1**). The impact of blur is particularly pronounced for high spatial frequency (SF) signals, hindering the perception of fine spatial details and sharp edges present in natural images. There is growing evidence that neural adaptation—a ubiquitous coding strategy of the brain—recalibrates visual processing in order to compensate for the eye’s optics, providing humans with a seemingly exceptional perceptual quality despite the poor quality of retinal images (Artal et al., 2004; Radhakrishnan et al., 2015; Sawides et al., 2011b; Webster, 2017, 2015; Webster et al., 2002). Vision is adapted to the natural level of blur present in the retinal image of each individual (Barbot et al., 2021; Ng et al., 2022, 2021; Sabesan and Yoon, 2009; Sawides et al., 2011b), with vision being the clearest with the eye’s own aberration pattern rather than with unfamiliar ones (Artal et al., 2004; Sabesan and Yoon, 2010; Sawides et al., 2011b). Neural adaptation to the eye’s optics affects contrast sensitivity (Barbot et al., 2021; Georgeson and Sullivan, 1975; Mon-Williams et al., 1998; Ng et al., 2022; Rajeev and Metha, 2010; Rouger et al., 2010; Sabesan and Yoon, 2009; Venkataraman et al., 2015), acuity (Barbot et al., 2021; Mon-Williams et al., 1998; Rosenfield et al., 2004; Rouger et al., 2010; Sabesan and Yoon, 2009), and perceived image focus (Artal et al., 2004; Sawides et al., 2011a, 2011b, 2010; Webster et al., 2002). However, the neural compensatory mechanisms underlying these adjustments remain unclear.

**Figure 1.**
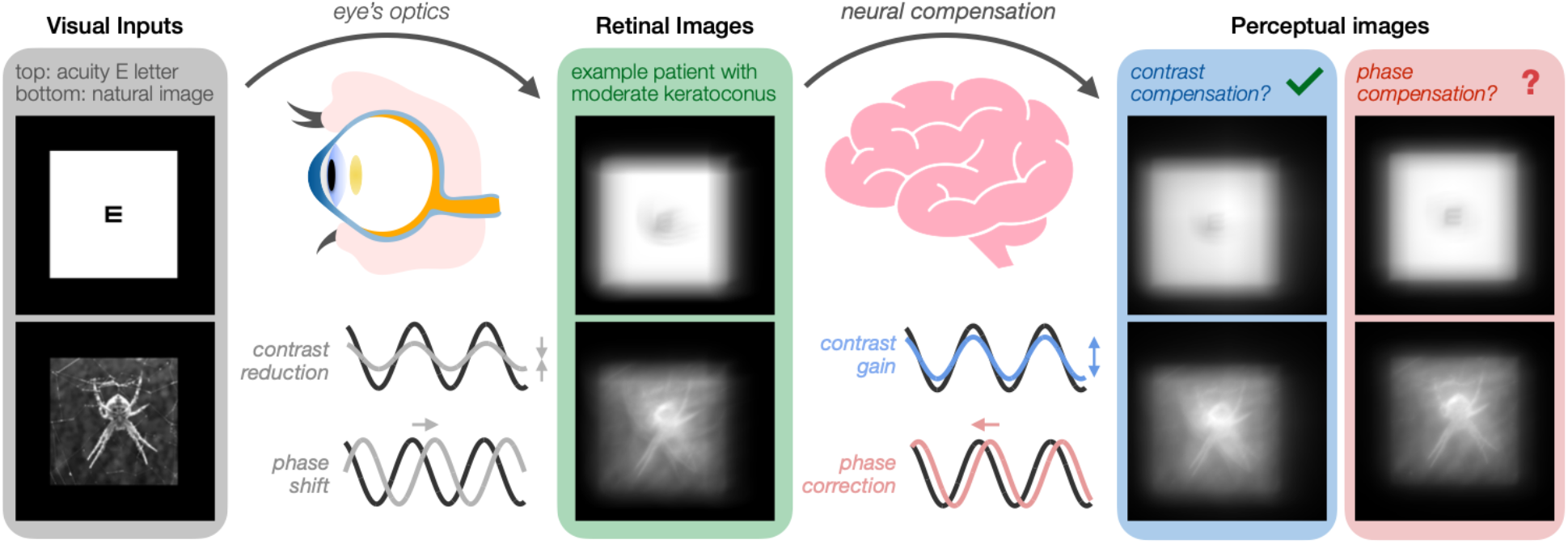
Neural adaptation to the eye’s optics. The eye’s optics degrade visual inputs transmitted to the retina in two ways: it reduces image contrast and can disrupt the phase (position) of visual signals relative to each other. Although there is evidence that neural adaptation to optical blur helps partially counteract blur-induced contrast reductions, it is unknown whether the visual system can compensate for phase disruptions. Given the importance of phase information in visual recognition and image quality, phase compensation may play a vital role in the brain’s ability to adapt to the eye’s optics to improve perceptual quality. *Left panel:* Visual inputs corresponding to a high-contrast acuity letter (20/20) and a grayscale image of a spider. *Middle panel:* Simulated retinal images using the habitual optical quality of a patient with moderate keratoconus–a corneal disease causing severe optical aberrations. *Right panels*: Simulated images illustrating the benefits of contrast compensation and phase compensation, respectively. Both mechanisms would be needed to optimally enhance perceived image quality.

One mechanism by which the visual system is able to attenuate the impact of the eye’s optics is through contrast compensation (**Fig.1**). Previous studies have provided evidence of contrast gain control mechanisms that help compensate for blur-induced reductions of high-SF signals in the retinal image (Barbot et al., 2021; Georgeson and Sullivan, 1975; Mon-Williams et al., 1998; Ng et al., 2022; Radhakrishnan et al., 2015; Rajeev and Metha, 2010; Venkataraman et al., 2015). Most of these studies have focused on lower-order optical aberrations of the eye (Georgeson and Sullivan, 1975; Mon-Williams et al., 1998; Rajeev and Metha, 2010; Rosenfield et al., 2004; Sawides et al., 2010; Venkataraman et al., 2015; Webster et al., 2002), such as defocus and astigmatism, which reduce image contrast without having a substantial impact on phase congruency; i.e., lower-order aberrations cause phase reversals but do not result in phase shifts per se. Contrast compensation mechanisms can therefore counteract the impact of relatively simple optical blur induced by lower-order aberrations, improving visual processing and perceptual quality over time.

However, optical quality is also constrained by higher-order aberrations (HOAs). While lower-order aberrations can be easily corrected using spectacles, contact lenses, or refractive surgery, HOAs are difficult to correct and limit visual processing and image quality, even in typical eyes (Liang et al., 1997; Sabesan et al., 2007; Yoon et al., 2004; Yoon and Williams, 2002). Importantly, HOAs not only reduce contrast but also disrupt the phase congruency of retinal images, which dramatically hinders perception. Indeed, whereas contrast is fundamental for the brain’s coding of visual information, perceptually-important information about recognizable features of natural images is almost exclusively contained within the phase spectrum of the image (**Fig.2**). Key structural features such as broadband edges correspond to locations of maximal local phase congruency (Henriksson et al., 2009; Kovesi, 2000; Morrone and Burr, 1988; Oppenheim and Lim, 1981; Wang and Simoncelli, 2004; Wichmann et al., 2006). Moreover, disruptions of phase congruency can affect the benefits of contrast compensation on perceptual image quality (Bex et al., 2009). Yet, it remains unknown whether the brain can counteract the phase scrambling effects caused by the eye’s optics. This knowledge gap is substantial given the role of phase information in visual recognition and image quality.

**Figure 2.**
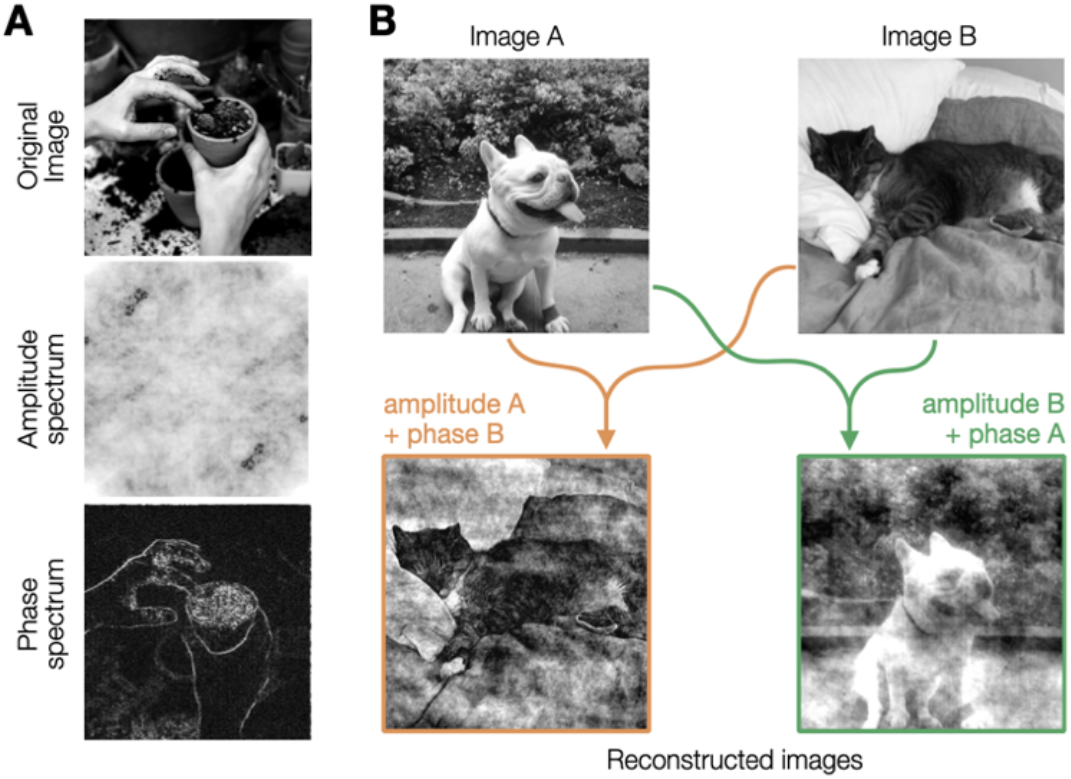
Perceptual importance of phase information. **(A)** Any image can be decomposed into its amplitude spectrum and phase spectrum. The phase spectrum, rather than the amplitude spectrum, contains most of perceptually-relevant information about image structure and is vital for visual recognition. **(B)** Reconstructed images reflect the phase spectrum, rather than the amplitude spectrum, of original images.

Here, we ask whether the visual system can compensate for the impact of optical aberrations via neural adaptation to the phase spectra. Such a phase compensation mechanism could play a vital role in the visual system’s ability to effectively attenuate the effects of blur on retinal image quality (**Fig.1**). Answering this question will provide key insights regarding how the brain adapts to the eye’s optics in order to help partially restore visual processing and perceptual quality over time.

Although HOAs remain relatively low in magnitude in typical eyes, they are usually uncorrected, limiting visual processing and perceptual quality (Artal et al., 2004; Liang et al., 1997; Yoon and Williams, 2002). HOAs can significantly increase with age (Amano et al., 2004), corneal surgeries (McCormick et al., 2005), or with corneal pathologies (Pantanelli et al., 2007). In such cases, the visual system is chronically exposed to substantial magnitudes of uncorrected optical aberrations, which severely degrade retinal image quality and impact visual processing. For instance, we previously showed evidence of altered contrast sensitivity and visual acuity following long-term exposure to severe optical aberrations in patients with keratoconus (KC)(Barbot et al., 2021; Ng et al., 2022; Sabesan and Yoon, 2009)–a corneal disease resulting in severe amounts of uncorrected HOAs. Notably, some of our previous findings suggest that long-term neural adaptation to the eye’s optics does not only affect contrast processing, but may also alter phase perception. Indeed, KC patients tested with their own native aberrations show better performance for broadband visual information (acuity letter), but no clear advantage for narrowband information (gratings), compared to typical observers tested with the KC patient’s own aberrations (Rouger et al., 2010; Sabesan and Yoon, 2010). While improved acuity with blur adaptation is often linked to enhanced contrast sensitivity of high SFs (Barbot et al., 2021; Rosenfield et al., 2004; Sabesan and Yoon, 2009), a phase compensation mechanism could also account for improved acuity. We hypothesize that the advantage for broadband information in KC patients might arise from phase compensation mechanisms that alter visual processing in order to counteract disruptions of phase congruency caused by uncorrected optical aberrations.

In the present study, we used adaptive optics (AO) (**Fig.3A**) to assess the impact of exposure to optical aberrations on perceived phase, from brief exposure to long-term adaptation. AO is a powerful technique that ensures precise, real-time control of optical quality during visual testing (Roorda, 2011), allowing to bypass the limit imposed by the eye’s optics to directly characterize changes in neural functions (Artal et al., 2004; Barbot et al., 2021; Liang et al., 1997; Ng et al., 2021; Sabesan and Yoon, 2009; Yoon and Williams, 2002). AO offers the unique possibility to fully correct all optical aberrations in both humans with typical eyes and patients with severe optical defects, as well as to induce sustained amounts of specific optical aberrations in order to assess how it affects visual functions. Correcting HOAs improves contrast sensitivity and acuity by enabling the detection of high-SF signals, which are otherwise indiscernible while viewing through the eye’s best refraction (Liang et al., 1997; Yoon and Williams, 2002). Moreover, the use of AO correction has revealed changes in contrast processing even for low SFs (Barbot et al., 2021; Ng et al., 2022), indicating that blur adaptation can have a broad impact on visual processing.

**Figure 3.**
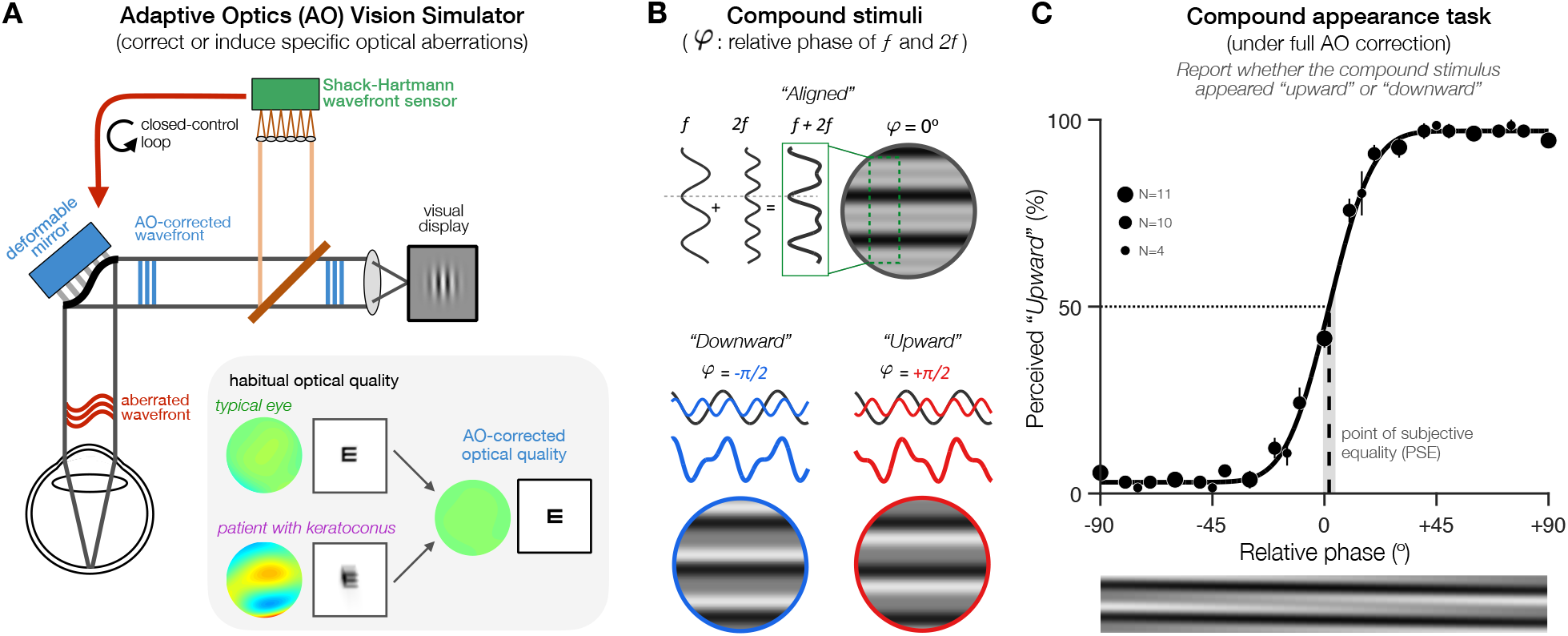
Experimental approach. **(A)** Schematic illustration of the AO vision simulator. AO correction allows us to bypass optical factors and compare visual functions under similar, fully-corrected optical quality in participants otherwise exposed to varying amounts of optical aberrations. Optical aberrations are measured in real-time by a Shack-Hartmann wavefront sensor. During closed-control loop, a deformable mirror maintains a desired aberration pattern by compensating for the participant’s wavefront aberrations. Wavefront maps and simulated acuity letter acuity are showed for participants with typical optical quality and moderate keratoconus. **(B)** Compound stimuli consisted of two suprathreshold horizontal gratings with frequencies *f* and *2f*. The appearance of compound stimuli depends on the *f*-*2f* relative phase, going from squarewave-like (*upper panel*) to sawtooth-like stimuli whose luminance peaks points either more “*downward*” (*lower left*) or more “*upward*” (*lower right*). **(C)** Perceived phase under full AO-correction in participants with typical optical quality (Exp.3, N=11). Perceived phase was measured by computing the percent of *“upward”* responses as a function of relative phase, and estimating the point-of-subjective equality (PSE)–i.e., the relative phase at which stimuli appeared aligned. Data were fit with a Cumulative Normal function to compute the PSE (+1.89º; vertical dashed line) with bootstrapped 95%-confidence intervals (95%-CI: [-0.7º +4.4º]; shaded gray area). All participants were tested across ±90º in relative phase, but the exact range of values used across participants in Exp.3 slightly varied (see *Methods*), as indicated by the size of each data point in this example figure. Error bars correspond to ±1SEM.

In a series of experiments, we assessed the impact of brief, short-term (∼1h), and long-term (months to years) exposure to specific optical aberrations on phase perception. We estimated perceived phase by asking participants to judge compound grating stimuli whose appearance depends directly on the phase alignment between two sinusoidal components (**Fig.3B,C**). Compound gratings are simplified broadband stimuli that have been successfully used to manipulate and study phase perception (Atkinson and Campbell, 1974; Badcock, 1984; Bennett and Banks, 1987; Burr, 1980; Henriksson et al., 2009; Mechler et al., 2002). First, we measured the effects of brief exposure to specific amounts of AO-induced vertical coma–a HOA resulting in disrupted phase congruency–on perceived phase (Exp.1). Then, we assessed changes in perceived phase before, during and following short-term (∼1h) exposure to AO-induced vertical coma (Exp.2). Finally, we investigated the impact of long-term adaptation to poor optical quality on phase perception in KC patients, who are chronically exposed to substantial amounts of uncorrected HOAs and disrupted phase in their daily life. Altogether, our results support the existence of neural adaptation mechanisms that can detect and compensate for blur-induced disruptions in phase congruency of the retinal image, adjusting phase perception to help improve perceptual quality over time.

## Results

In all experiments, we asked participants to judge compound gratings whose appearance depends on the relative phase between the two frequency components *f* and *2f* (**Fig.3B**). At the reference point (relative phase: 0º), the troughs of *f* and *2f* sinusoids are aligned and compound stimuli appear ‘*square-wave-like’*. Varying the relative phase between *f* and *2f* results in ‘*saw-tooth-like*’ compound gratings pointing either ‘*downward’* (−90º) or ‘*upward’* (+90º). Note that the reference phase of the *f* component varied randomly on each trial to avoid low-level effects related to local luminance contrast. We computed the proportion of trials in which the stimulus was perceived ‘*upward*’ as a function of relative phase, and fitted the data to estimate the point-of-subjective equality (PSE)–i.e., the relative phase at which compound patterns are perceived neither ‘*upward’* or ‘*downward’* (**Fig.3C**). Under full AO correction, the PSE at which compound patterns appear ‘*aligned’* corresponded to a relative phase of ∼0º, as expected (**Fig.3C**). We first validated our experimental approach by showing that physically shifting the relative phase by specific offsets resulted in PSE shifts whose magnitude and direction matched the offset added (***Fig*.*S1*** and *Supp. Information*). We then used this compound appearance task to assess changes in perceived phase at various timepoints: with brief exposure (Exp.1), during short-term adaptation (Exp.2), and following long-term exposure (Exp.3) to optical aberrations.

### Exp.1 – Perceived phase shifts with brief exposure match optical theory predictions

In Exp.1, we measured shifts in perceived phase occurring with brief exposure to specific amounts of vertical coma (**Fig.4**). We used AO to correct all optical aberrations and induce vertical coma–a HOA resulting in asymmetric blur that specifically affects horizontally-oriented spatial information. We computed the expected contrast reduction and phase shift in the presence of specific amounts of vertical coma from optical theory (see *Methods*). Specifically, contrast reduction and phase disruption are characterized by the Modulation Transfer Function (MTF) and the Phase Transfer Function (PTF), respectively. First, we predicted shifts in perceived phase (PSEs) for various amounts of vertical coma by simulating retinal images using Fourier propagation (**Fig.4B**). As expected, larger shifts in relative phase were predicted for larger magnitudes of vertical coma and for higher SFs, which were in opposite directions for positive and negative vertical coma (**Fig.4B**). We selected a range of vertical coma values for testing (±0.5µm, for *f*=3 cycles/deg.) that could be effectively evaluated using the compound appearance task (see *Methods*). We validated the predictions for these conditions directly from the AO visual display by capturing images of horizontal gratings under various amounts of AO-induced vertical coma. The phase shifts we obtained from the display matched the predictions from optical theory (**Fig.4C**).

**Figure 4.**
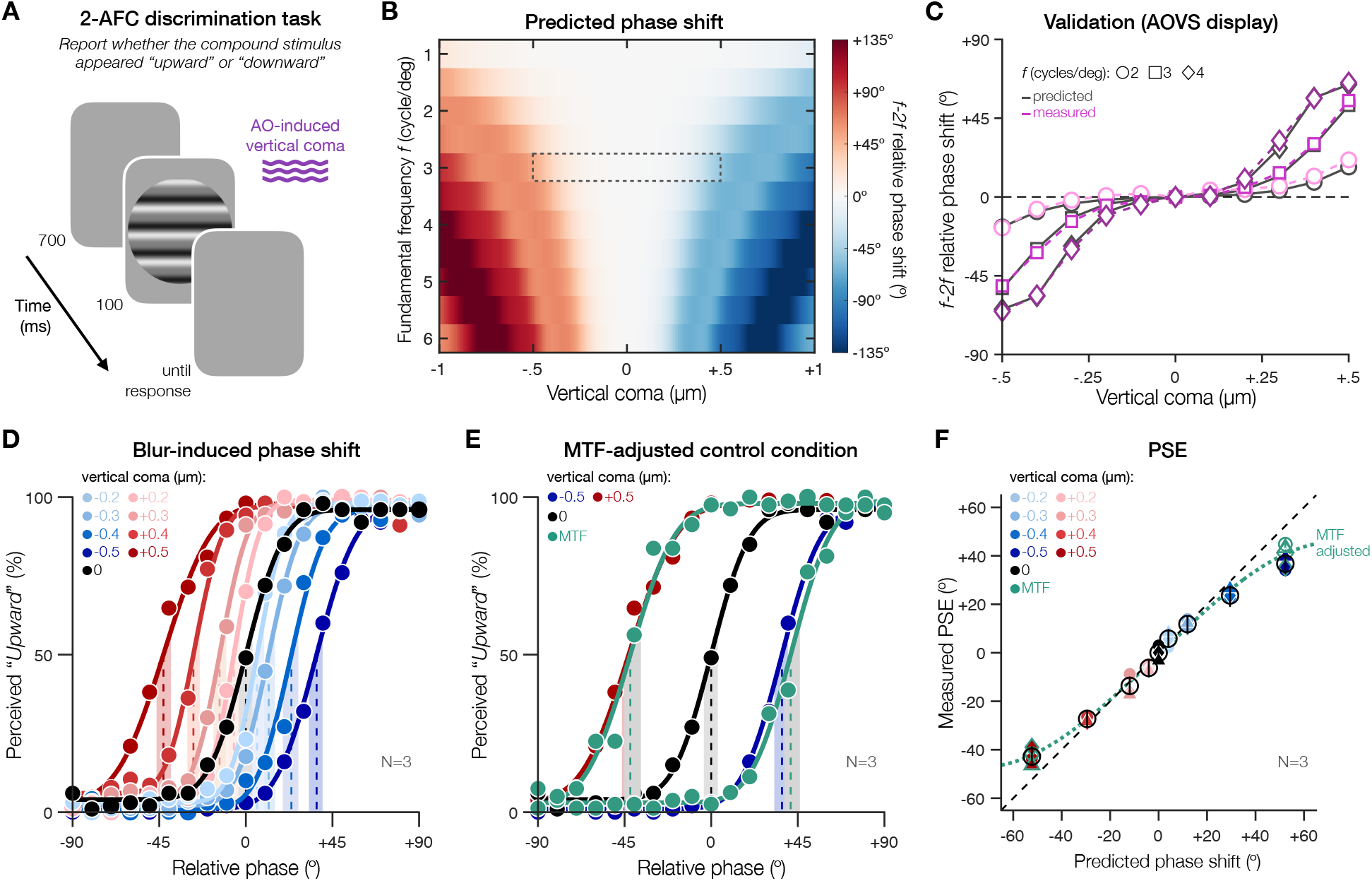
Perceived phase shifts with brief exposure to vertical coma. **(A)** *Trial sequence*. Participants judged the appearance of compound gratings under specific amounts of AO-induced vertical coma. **(B)** *Optical theory predictions*. We predicted the magnitude and direction of *f-2f* relative phase shifts for varying amounts of coma and different *f* frequencies. Dashed area indicates conditions selected for testing in (D-F). **(C)** *Setup validation*. We validated the predictions before testing directly from the visual display by capturing images of gratings under AO-induced vertical coma. **(D)** *Perceived phase shifts*. Brief exposure to vertical coma resulted in PSE shifts matching optical theory predictions, except at ±0.5µm. **(E)** *Impact of reduced contrast on perceived phase*. The attenuation of PSE shifts under ±0.5µm of coma was replicated under AO correction using MTF-adjusted stimuli with ±52.3º phase offset. **(F)** Perceived PSE estimates plotted relative to optical theory predictions. Individual PSEs correspond to filled colored symbols, while open black circles show group-average PSEs with 95%-CI error bars. As showed in (F), individual participant data were consistent with group-average data plotted in (D,E). Results in (D,E,F) are for compound gratings with a fundamental frequency of 3 cycles/deg (see **Fig.S2** for additional results using *f*: 4 cycles/deg).

Three participants with typical eyes judged compound grating stimuli (*f*: 3 cycles/deg) under various amounts of AO-induced vertical coma (**Fig.4D-F**). Under full AO correction, PSE estimates were centered on 0º (**Fig.4D**; -0.003º [-3.8º +3.4º]). In the presence of vertical coma, participants experienced shifts in perceived phase that matched optical theory predictions for mild-to-moderate amounts of vertical coma, both in terms of the magnitude and direction of the shift (**Fig.4D,F**). However, PSE shifts were attenuated compared to optical theory predictions and AO display validation for ±0.5µm of vertical coma (predicted: ±52.3º; perceived: -42.82º [-38.9º -46.4º] and +36.77º [+32.8º +39.9º], for +0.5 and -0.5 µm, respectively) (**Fig.4D-F**).

What could explain the mismatch between observed and expected phase shifts for larger amounts of vertical coma? We know that we can measure PSE shifts up to ±60º using high-contrast stimuli (**Fig.S1**). Moreover, we validated that there was no phase deviation in our AO display that could explain the deviations observed with perceived phase shifts (**Fig.4C**). We hypothesized that this mismatch might therefore reflect the impact of blur-induced contrast reduction on perceived phase. Although we maintained the contrast ratio between the *f* and *2f* components in the presence of vertical coma (see *Methods*), reductions in contrast at ±0.5µm of vertical coma might be too severe for the *2f* component and affect the appearance of compound gratings, thus attenuating the magnitude of PSE shifts.

We directly tested this hypothesis by measuring PSEs under AO correction using compound stimuli physically-adjusted both in terms of the amplitude and phase of the *f* and *2f* components (i.e., using MTF and PTF values for ±0.5µm of vertical coma). Relative to the actual offset added to these MTF-adjusted stimuli (±52.3º), participants showed attenuated PSE shifts (**Fig.4E**; -42.02º [-36.5º -45.1º] and +41.3º [+38º +46.1º], for ±52.3º phase offset) that matched with the phase shifts observed under ±0.5µm of vertical coma (−42.82º [-38.9º -46.4º] and +36.77º [+32.8º +39.9º] for +0.5 and -0.5µm, respectively). In other words, contrast reductions can affect the magnitude of perceived phase shifts. The results were consistent across individual participants (see **Fig.4F** for individual and group average PSE estimates). Furthermore, using higher-SF stimuli (*f*: 4 cycles/deg; **Fig.S2**) led to a stronger attenuation of PSE shifts, which was observed for vertical coma values above 0.3µm. These findings show that brief exposure to vertical coma results in perceived phase shifts that match predictions from optical theory but can be attenuated by contrast reductions.

### Exp.2 – Short-term adaptation to vertical coma compensates for phase disruptions

In Exp.2, we assessed whether the visual system can adaptively compensate for the impact of vertical coma on phase perception over relatively short periods of time (**Fig.5**). To do so, we measured PSEs before, during and following short-term (∼1h) exposure to -0.4µm of vertical coma. We selected this value based on Exp.1 to maximize our chance to observe adaptation effects while minimizing the impact of blur-induced contrast reductions. Each experimental session was divided into 3 periods during which participants continuously performed the phase appearance task (**Fig.5A**): baseline (∼8min), blur adaptation (∼70min), and post-adaptation (∼12min). Baseline and post-adaptation periods were similar: participants judged the appearance of compound gratings presented under AO correction (*PSE*_*predicted*_: 0º). During adaptation, participants were exposed to a sustained amount of AO-induced vertical coma (−0.4µm; *PSE*_*predicted*_: +29.5º). The first 3 adaptation blocks (∼3min) consisted of compound gratings only to accurately estimate the blur-induced PSE shift. Then, each compound grating was sandwiched between a series of high-contrast grayscale images consisting of complex natural images and checkerboard patterns (**Fig.5B,C**). These images appeared blurry owing to the presence of vertical coma, providing invariants the visual system can use to detect and adapt for the impact of blur (e.g., hard edges should have all SFs in sine phase). Adaptation stimuli encompassed a wide range of visual properties and varied rapidly to homogenize local light adaption and minimize afterimages (Elliott et al., 2011, 2007; Sawides et al., 2011a), ensuring adaptation to the presence of vertical coma rather than to specific image properties. To better assess PSE changes over time, we computed PSEs by binning trials from adjacent (±1) blocks, except if the trials belonged to different time periods (see *Methods*, and **Fig.S3** for results using unbinned data).

**Figure 5.**
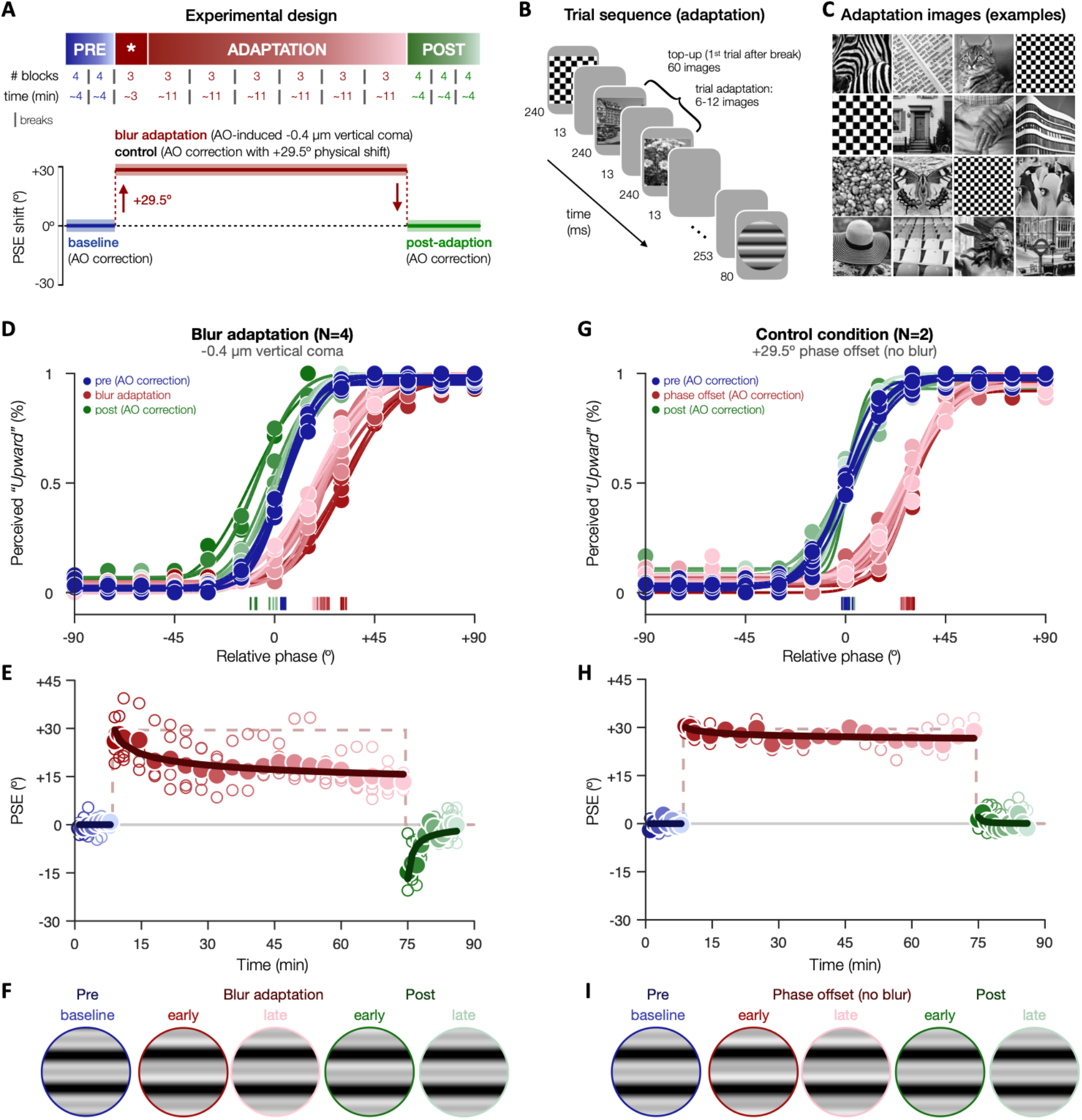
Short-term adaptation to vertical coma. **(A)** Each experimental session consisted of 41 blocks divided into 3 segments: baseline (pre-adaptation), adaptation, and post-adaptation. Both pre- and post-adaptation were measured under AO correction (*PSE*_*predicted*_:0º). During blur adaptation, -0.4µm of vertical coma was AO-induced (*PSE*_*predicted*_:+29.5º). After the first 3 blocks of adaptation (indicated by the * symbol), we started presenting grayscale natural images and checkerboards before each compound grating, serving as cues the visual system could use to detect and adapt to blur. In a control condition, AO correction was maintained during the entire session and a phase offset (+29.5º) was added instead, thus mimicking the impact of vertical coma on perceived phase without observers being exposed to blur. Breaks were allowed at specific time points, as indicated by vertical lines. **(B)** Trial sequence during the adaptation (and control) segment. **(C)** Examples of adaptation stimuli. **(D–F)** *Blur adaptation*. Before adaptation, psychometric functions and PSE estimates were centered near 0º. Blur exposure resulted in a PSE shift that initially matched optical theory predictions (+29.5º), but then decreased over time. As soon as blur was removed (post-adaptation), an aftereffect in the opposite direction was observed. **(G–I)** *Control experiment*. In the absence of blur, PSE shifts matched the added phase offset (+29.5º) and remained stable over time, with post-adaptation PSEs returning immediately to baseline. (D,G) Psychometric curves fitted to group-average data. (E,H) Group-average (filled circles) and individual (open circles) PSE estimates, with solid lines corresponding to power functions fitted to group-average PSEs. To better visualize changes in PSE over time, PSE estimates are adjusted for small biases observed during baseline, thus treating 0º as baseline. (F,I) Schematic representations of perceived phase over time observed in both experimental conditions.

Our results provide compelling evidence of neural adaptation to the phase spectra during short-term exposure to vertical coma (**Fig.5D-F**). Before blur exposure, PSEs were near 0º, with a slight positive bias on average during baseline (+3.93º±0.81º; **Fig.5D**). To treat 0º as reference point, we subtracted the mean of PSE estimates during baseline from all PSE estimates (**Fig.5E**). Inducing vertical coma resulted in a significant PSE shift relative to 0º baseline (+26.04º [+18º +33.8º]), which matched the prediction from optical theory (*PSE*_*predicted*_: +29.5º). Critically, the magnitude of the blur-induced phase shift decreased during blur exposure, by -12.7º by the end of the adaptation period (last block: +13.34º [+6.2º +20.8º]). This attenuation over time was well captured by a power function (*R*^2^_adj_ =.77, *p*<.001), corresponding to ∼47% reduction by the end of adaptation. In addition, when we removed AO-induced vertical coma (post-adaptation), PSEs did not return to baseline levels immediately. Instead, an aftereffect was observed in the opposite direction (−14.72º [-21.8º -8.0º]), which represents a hallmark of visual adaptation. PSEs returned to baseline level after ∼3-4min (5^th^ post-adaptation block: -3.18º [-8.8º +2.7º]). Importantly, this pattern of results did not depend on whether PSE estimates were obtained from binned (**Fig.5**) or unbinned data (**Fig.S3**). All participants showed an attenuation of PSE shifts during adaptation (by -12.1º±6.4º on average; **Fig.S3**), with 3 out of 4 participants showing an aftereffect (−13.3±7.6º).

In a control condition, we ruled out confounding factors such as changes in observers’ response bias over time due to fatigue or serial dependency effects, as well as low-level effects of the adaptation stimuli presented during the adaptation period. To do so, we maintained AO correction during the entire control experiment, thus removing any blur-induced phase disruptions the visual system could adapt to. Instead, during the adaptation period, a phase offset of similar magnitude and direction than with -0.4µm of vertical coma (+29.5°) was added to compound stimuli. In the absence of vertical coma, adaptation stimuli should be perceived in-focus and should not induce blur adaptation, whereas the phase offset should result in a change in the observer’s response pattern. As expected, we initially observed a significant PSE shift relative to baseline that matched the phase offset added to the stimuli (+30.4º [+23.4º +37.7º]), matching the added phase offset (**Fig.5G-I**). However, the PSE shift remained stable during the entire exposure period (last block: 28.86º [+21.7º +35.3º]), returning immediately to baseline once the phase offset was removed (+1.36º [-5.82º +8.1º]). This pattern was clear across both participants (**Fig.S4**). Thus, the attenuation of perceived phase shifts during short-term blur exposure, and the subsequent aftereffect, likely reflect neural adaptation to phase spectra in the presence of vertical coma, rather than non-perceptual changes in response bias or low-level interactions during blur adaptation.

### Exp.3 – Altered phase perception following long-term exposure to poor optics optical aberrations

Finally, we assessed the effects of long-term exposure (months to years) to optical aberrations on phase perception. To do so, we measured perceived phase in patients with keratoconus (KC)(**Fig.6A**). KC patients are chronically exposed to large amounts of uncorrected HOAs in their daily life, with particularly large amounts of vertical coma. Long-term exposure to poor optical quality induces pronounced neural compensatory adjustments in KC patients. For instance, when tested under full AO correction, KC patients show poorer acuity and altered contrast sensitivity than healthy participants tested under similar, AO-corrected optical conditions (Barbot et al., 2021; Hastings et al., 2021; Ng et al., 2022; Rouger et al., 2010; Sabesan et al., 2017; Sabesan and Yoon, 2009). Here, we tested whether KC patients show evidence of altered phase perception under fully-corrected optical quality, which would suggest long-term neural compensatory adjustments to phase spectra.

**Figure 6.**
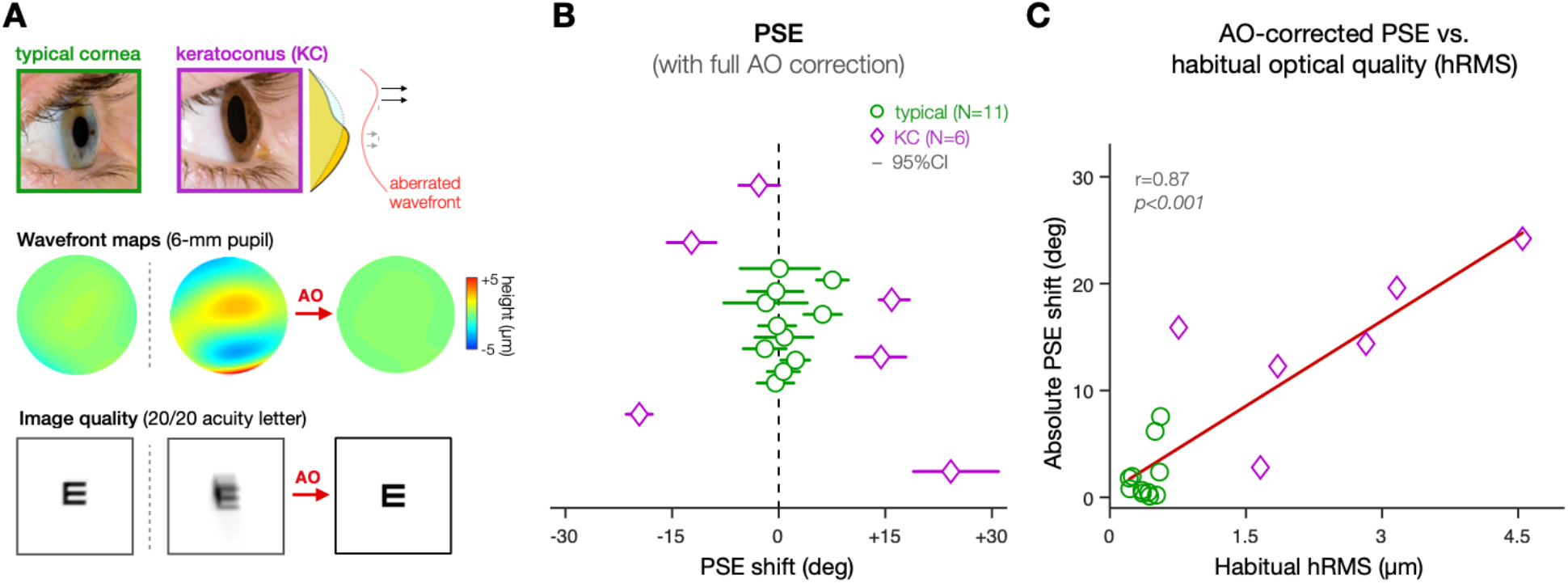
Altered phase congruency following long-term adaptation to poor optical quality. **(A)** Relative to typical ‘healthy’ eyes, KC eyes are affected by large amounts of optical aberrations, as illustrated by the wavefront maps. As a result, the visual system of KC patients is constantly exposed to degraded retinal images, as illustrated by simulated acuity letters. AO allows to fully correct all optical aberrations while assessing visual functions, even in KC eyes. **(B)** Under full AO correction, participants with healthy eyes showed PSEs tightly distributed around 0º, as expected. In contrast, KC patients showed larger PSE shifts, consistent with altered phase congruency. Each data point corresponds to PSE estimates (x-axis) of individual participants (y-axis) plotted with 95%-CIs (see also Fig.S5). **(C)** The magnitude of the PSE shift correlated with the amount of habitual optical aberrations (e.g., higher-order aberrations–hRMS) that each participant is exposed to in their daily life (see also Fig.S6).

To do so, we measured perceived phase under AO correction in participants with typical eyes (N=11) and patients with mild-to-severe KC (N=6)(**Fig.S5**). For each participant, we estimated the habitual optical quality as the total RMS error (tRMS) and higher-order aberration RMS error (hRMS) obtained from wavefront measurements (see *Methods*). Relative to participants with typical eyes (tRMS: 1.06±0.31µm; hRMS: 0.4±0.13µm), KC patients were exposed to poorer retinal image quality in their daily life (tRMS: 4.04±2.32µm; hRMS: 2.47±1.33µm). As shown in previous studies(Barbot et al., 2021; Hastings et al., 2021; Ng et al., 2022; Rouger et al., 2010; Sabesan et al., 2017; Sabesan and Yoon, 2009), AO correction allowed us to bypass optical factors and maintain total RMS around 0.05µm, with no difference between typical (0.051±0.01) and KC (0.058±0.01) eyes (Welch’s t-test: t=1.22, *p=*.*260*, Cohen d=0.66) (**Fig.S6A**). This allowed us to measure PSE estimates for each participant under similar, AO-corrected optical quality (**Fig.6** and **Fig.S5**).

Under fully-corrected optical quality, participants with healthy eyes had PSEs distributed around 0º (**Fig.6B**; +1.17º±3.07º; Wilcoxon test: V=42, *p=0*.*465*, d=0.27). In contrast, KC patients exhibited clear shifts in perceived phase away from 0º under similar AO-corrected optical quality (**Fig.6B**), resulting in larger variance in the distribution of PSE estimates between the two groups (Levene’s test for equality of variance: F=39.17, *p*<0.001). Relative to the magnitude of absolute PSE shifts observed in participants with typical eyes (2.04º±2.52º), KC eyes exhibited significantly larger PSE shifts (14.86º±7.26º)(Welch’s t-test: t=4.19, *p=*.*007*, Cohen d=2.36). It is worth re-emphasizing that both control participants and KC patients were tested under similar aberration-free optical quality, for which PSEs are expected to remain around 0º. Moreover, we found no correlation between the magnitude of individual PSE shifts and the amount of residual aberrations under AO correction (**Fig.S6A**; r=.26, *p*=.31). Thus, the difference in PSE we observed likely reflects the impact of neural compensatory adjustments on phase processing following long-term exposure to optical aberrations. Consistent with this idea, the magnitude of individual PSE shifts correlated with the amount of habitual optical aberrations experienced by each participant in their daily life (**Fig.6C**–hRMS: r=.87, *p*<.001; **Fig.S6B**–tRMS: r=.82, *p*<.001). Finally, larger PSE shifts under AO correction were associated with stronger deficits in high-contrast letter acuity under full AO correction (**Fig.S6C**; second-order polynomial: F=23.3, *p*<.001, *R*^*2*^_*adj*_ *= 0*.*74*). These results provide evidence that long-term exposure to severe optical aberrations alters perceived phase in KC patients, consistent with the existence of neural compensation adjustments to phase spectra.

## Discussion

The present study provides vital insights into comprehending how the brain’s coding of sensory information adapts to the structure of the degraded retinal image to help provide a clearer and sharper perceptual experience. AO provided full control over optical factors, allowing us to efficiently induce specific amounts of vertical coma (Exp.1,2) and to test human observers under fully corrected optical conditions even in severely aberrated KC eyes (Exp.3). Our results unveil the visual system’s capacity to adapt visual processing to the phase spectra of the retinal input in order to counterbalance the impact of optical aberrations on phase congruency. We substantiated the existence of this novel compensatory mechanism with evidence of both short-term and long-term neural adaptation effects to the eye’s optics on perceived phase.

A key function of human perceptual systems is to adapt to the environment. Human vision readily detects blurring of retinal images, and makes compensatory adjustments to render the retinal image clearer (Artal et al., 2004; Georgeson and Sullivan, 1975; Sawides et al., 2011b; Webster et al., 2002). Several studies have showed that the visual system relies on contrast gain control mechanisms to compensate for reductions in image contrast induced by blur (Barbot et al., 2021; Georgeson and Sullivan, 1975; Mon-Williams et al., 1998; Ng et al., 2022; Rajeev and Metha, 2010; Venkataraman et al., 2015; Webster, 2017, 2015). Such a compensation mechanism is vital given the significant impact contrast information has on visual performance and image quality. Nevertheless, perceptually-relevant information about recognizable features of natural images resides almost exclusively within the phase spectrum (Henriksson et al., 2009; Kovesi, 2000; Morrone and Burr, 1988; Oppenheim and Lim, 1981; Wang and Simoncelli, 2004; Wichmann et al., 2006). The visual system depends on phase information to detect edges and boundaries, which are fundamental for image segmentation and visual recognition. Although we did not assess performance and perceived focus per se, it is reasonable to infer that phase compensation would help improve visual processing and perceptual quality in the presence of blur. In addition, we found that contrast reductions can affect perceived phase, which may impact neural adaptation to the phase spectra. Moreover, contrast sensitivity (Bex et al., 2009) as well as perceived blur (Murray and Bex, 2010) depends on the relative phase of natural images. These findings suggest that contrast gain control and phase compensation are two neural adaptation mechanisms that likely interact with each other, allowing visual processing to recalibrate itself to the natural level of blur present in the retinal image of each individual (Barbot et al., 2021; Ng et al., 2022, 2021; Sabesan and Yoon, 2010, 2009; Sawides et al., 2011a, 2011b)

The visual system’s ability to compensate for phase disruptions caused by the eye’s optics suggests that the brain possesses a representation of what constitutes an optimal (i.e., unaberrated) image. Invariants present in natural scenes play a key role by enabling the visual system to detect and correct for the presence of blur. Phase relations are inherently robust in the optical images of objects present in our natural environment, with edges corresponding to points of maximum local phase congruency (Henriksson et al., 2009; Kovesi, 2000; Morrone and Burr, 1988; Oppenheim and Lim, 1981; Wang and Simoncelli, 2004). Phase coding in the visual system is ‘tuned’ to those phases, with infants developing sensitivity to spatial phases by 2-3 months of age (Braddick et al., 1986). Natural images, as used in our study, are typically chosen to induce blur adaptation and evaluate perceived image focus (Radhakrishnan et al., 2015; Sawides et al., 2011b, 2010; Venkataraman et al., 2015; Webster et al., 2002), as they contain invariants the visual system relies on for signaling the presence of blur and recalibrating itself.

The present study helps reconcile earlier observations of long-term neural compensation in KC patients. As previously mentioned, when tested under their own native optical aberrations, KC patients exhibit better letter acuity than typical observers tested under similar KC’s optical aberrations, all while showing no clear advantage in contrast sensitivity (Rouger et al., 2010; Sabesan and Yoon, 2010). Some authors have interpreted these findings as evidence that neural adaptation to poor retinal image quality depends on the visual task (Rouger et al., 2010), occurring for real-life tasks such as letter acuity tasks but not with grating stimuli. It is also important to note that visual acuity measurements alone do not allow to differentiate between contrast gain adjustments and phase compensation, as both mechanisms would improve broadband acuity. Our results provide evidence of poorer letter acuity and altered perceived phase in KC patients under AO correction, with the magnitude of both effects being positively correlated to the amount of habitual optical aberrations. Note that KC patients exhibit neural insensitivity for higher SFs relative to typical eyes under AO correction (Barbot et al., 2021; Hastings et al., 2021; Ng et al., 2022; Sabesan et al., 2017; Sabesan and Yoon, 2009), which could have attenuated the magnitude of the shifts in perceived phase we observed. The present findings support the hypothesis that long-term exposure to poor optical quality in KC alters phase processing, which would improve the processing of broadband information, such as acuity letters and natural images, without affecting narrowband Gabor stimuli typically used in contrast sensitivity tasks. Given that neural compensation to the eye’s optics remain limited and perceptual quality poor in KC patients, our results hold promise for developing better clinical care and visual rehabilitation strategies.

Although the present results hold significance, we should be careful not to overgeneralize our findings. Compound gratings are simplified broadband stimuli that have been successfully used to study phase perception (Atkinson and Campbell, 1974; Badcock, 1984; Bennett and Banks, 1987; Burr, 1980; Henriksson et al., 2009; Mechler et al., 2002), allowing us to efficiently manipulate and predict phase shifts for specific amounts of vertical coma. While the power of our approach lies in its simplicity, the experimental conditions were optimized to potentially induce neural adaptation to the phase spectra; i.e., by testing the impact of vertical coma on horizontal compound gratings. As such, our results may not fully capture the complexity of neural adaptation to the eye’s optics. Natural images encompass a wide spectrum of orientations and SFs, which likely interact and affect blur adaptation. Predicting perceived phase shifts becomes progressively intricate when using more natural stimuli, or when considering more severe habitual aberration patterns, such as in KC eyes. For instance, while larger amounts of habitual optical aberrations were associated with larger shifts in perceived phase in KC patients, we observed PSE shifts in both directions that cannot simply be accounted by the amounts of vertical coma present in KC eyes. Moreover, it’s important to note that most optical aberration patterns do not necessarily lead to phase disruptions. Thus, there remains a need for further investigation into the broader, overall contribution of phase compensation mechanisms to neural adaptation to the eye’s optics.

Our results are largely agnostic regarding the neural bases by which the visual system detects disruptions in local phase congruence and recalibrates sensory processing accordingly. The visual system is sensitive to differences in relative phase between SF components, a trait observed as early as the primary visual cortex (V1)(Aronov et al., 2003; Henriksson et al., 2009; Mechler et al., 2002; Williams and Shapley, 2007). Phase-sensitive pooling of SF information represents a statistically optimal way to analyze the output from V1 neurons (Hyvarinen et al., 2005). In the macaque V1, phase-selective mechanisms, as well as to phase differences in compound grating stimuli (Henriksson et al., 2009). However, selectivity for phase congruency increases along the visual hierarchy (Henriksson et al., 2009), and only higher-level areas can discriminate between broadband edges and lines (Perna et al., 2008). These findings support the idea that higher visual areas mediate phase-sensitive pooling of SF information across multiple spatial scales, which is required for human perception of broadband natural images (Henriksson et al., 2009). The detection of disrupted local phase congruency in natural images likely engages higher-level areas that, through feedback, modulate phase-sensitive mechanisms in early visual regions.

While a single neural adaptation mechanism operating over different timescales could account for both short-term and long-term adaptation effects, there is evidence of distinct adaptation mechanisms tuned to specific timescales (Bao and Engel, 2012; Webster, 2015), which may affect neural compensation to the eye’s optics in qualitatively different ways. Considering that phase-sensitive SF pooling is ubiquitous in the human visual cortex, short-term and long-term adaptation effects may reflect changes at different stages of visual processing. Moreover, while short-term adaptation generally yields relatively brief after-effects, readaptation to enhanced optics in patient populations often occurs over longer timeframes and may involve perceptual learning interventions (Sabesan et al., 2017; Vinas et al., 2012). Neural adaptation to phase spectra might also differ across the visual field, given differences in phase perception between central and peripheral visual locations (Bennett and Banks, 1987; Morrone et al., 1989). Pattern perception appears to rely on combined phase-amplitude mechanisms (Freeman and Simoncelli, 2011; Zhang et al., 2014), with a greater emphasis on phase information near the fovea and an increasing role of amplitude in the periphery. These results suggest that neural adaptation to the eye’s optics might be eccentricity-dependent, potentially resulting in weaker adaptation to phase spectra at peripheral locations.

Neural adaptation to the phase spectra should benefit binocular vision (Ng et al., 2021), particularly given the importance of phase information in binocular disparity (Anzai et al., 1997). It is worth emphasizing that our study focused on the effects of neural adaptation on perceived phase only in one eye. However, neural compensation for the eye’s optics likely involves both eyes. Under natural viewing, individuals are chronically exposed to interocular differences in blur, which potentially leads to distinct adjustments in each eye over extended timeframes. Evidence suggests that the retinal image quality of the eye with the best optics serves as reference, with perceived best focus measured independently in each eye matching the optical blur of the better eye(Kompaniez et al., 2013; Radhakrishnan et al., 2015). Future research should investigate the impact of interocular differences on phase perception and neural adaptation, over both short- and long-term time periods.

In summary, the present study reveals the existence of neural adaptation mechanisms that detect and mitigate signal disruptions in local phase congruency by recalibrating visual processing. Given the critical role of phase information in visual recognition and perceived image quality, neural adaptation to the phase spectra emerges as a potentially vital mechanism for enhancing perception in the presence of blur. These findings deepen our understanding of the adaptive mechanisms by which the brain optimizes perception despite the limitations of the human eye, offering valuable implications for fields spanning neuroscience, ophthalmology, and the broader study of neuroplasticity and visual rehabilitation.

## Methods

### Participants

All participants were cyclopleged with tropicamide (1%) to dilate their pupil and paralyze accommodation during testing, after being screened by one of our ophthalmologists to ensure dilation was safe and provide standard information (e.g., corneal curvature, refractive error). An additional screening session was performed to ensure we could obtain good quality wavefront measurements and reach stable AO correction. Habitual optical quality was estimated for each participant from wavefront measurements collected using their everyday correction method, if any. The square root of the sum of Zernike coefficients for a 6-mm pupil was used to estimate total RMS error (tRMS: all coefficients, except piston and prisms) and HOAs RMS error (hRMS: 6–65th coefficients). Three participants (3 males; 22-51yo) with typical optical quality (tRMS: 1.04µm, range: 0.57-1.42µm; hRMS: 0.44µm, range: 0.35-0.54µm; 6-mm pupil) completed Exp.1 and S1 (**Fig.4,S1**). Exp.2 included an additional participant (25yo female) with typical optical quality (RMS: 1.59µm, hRMS: 0.56µm). Exp.3 included eleven participants with typical eyes (8 males, 32±11.8yo; tRMS: 1.06±0.31µm; hRMS: 0.4±0.13µm) and six KC patients (5 males, 35.7±15.5yo; tRMS: 4.04±2.32µm; hRMS: 2.47±1.33µm). All control observers were emmetropes or low myopes/hyperopes with well-corrected optical quality in both eyes. All patients had been diagnosed with KC at least 12 months before testing (reported onset around their twenties), and had been wearing the same habitual corrective method without substantial changes in their vision for at least 6 months before testing. The eye with the most severe optical aberrations was selected for monocular testing as it is generally associated with stronger adaptation effects (Barbot et al., 2021; Ng et al., 2022; Sabesan et al., 2017; Sabesan and Yoon, 2009), except for one patient due to the presence of scarring in the worst eye incompatible with AO correction. Two KC patients and one control observer did not pass the screening stage, and were not tested further along. The Research Subjects Review Board at the University of Rochester approved all experiments, which adhered to the tenets of the Declaration of Helsinki and were conducted after obtaining informed, written consent. Participants were compensated$12/hr.

### Adaptive Optics Vision Simulator

Optical quality and visual performance were assessed under white light conditions using an Adaptive Optics Vision Simulator (AOVS). As previously detailed (Barbot et al., 2021; Ng et al., 2022, 2021; Sabesan et al., 2017; Sabesan and Yoon, 2010, 2009; Zheleznyak et al., 2016), our AOVS (**Fig.3A**) consists of a custom-built Shack–Hartmann wavefront sensor that measures observers’ wavefront aberrations, and a deformable mirror (ALPAO-DM97, Montbonnot, France) used to correct and/or induce wavefront aberrations online. An artificial pupil controls pupil size while visual stimuli are presented using a modified digital light processor (8bits, SharpXR-10X, Osaka, Japan). The visual display subtended 3.98×2.98deg (1024×768, 75Hz; ∼0.23 arc/min per pixel) and was gamma-corrected with a PR-650 SpectraScan Colorimeter (PhotoResearch, Chatsworth, CA). A superluminescent diode (840±20nm) was used for wavefront sensing. Head movements were stabilized with a headrest and a dental impression bite bar mounted to x-y-z translation stages. A camera imaging observers’ pupil was used to maintain pupil alignment. The AOVS operates in a continuous closed loop between wavefront sensing and deformable mirror correction, at ∼8Hz. Full correction of all coefficients was applied during testing, with the exception of defocus for which subjective values were used to correct axial chromatic aberrations and ensure the best focus in white light conditions. Subjective defocus was estimated by asking participants to adjust defocus using a motorized Badal prism while correcting all other aberrations in order to maximize image quality. We verified that subjective defocus values provided the highest acuity and perceived image focus under AO correction. Consistent with previous studies (Barbot et al., 2021; Ng et al., 2022, 2021; Sabesan et al., 2017; Sabesan and Yoon, 2010, 2009; Zheleznyak et al., 2016), the AOVS offered full control of wavefront aberrations over sustained periods (∼0.05µm residual RMS), in both ‘healthy’ and severely aberrated eyes.

### Compound grating stimuli

Suprathreshold horizontal compound stimuli were created by adding two sinusoids with frequency *f* and *2f*. The relative phase of the two components was shifted from a reference point at which the troughs of the two sinusoids are aligned and compound stimuli appear ‘*square-wave-like’* (0º relative phase)(**Fig.3B**). Varying the *f*-*2f* relative phase resulted in ‘*saw-tooth-like*’ stimuli pointing either ‘*downward’* (−90º) or ‘*upward’* (+90º). The absolute phase of the fundamental *f* was randomly set on each trial to avoid low-level repetition effects related to local image contrast. A raised-cosine envelope (2.6deg diameter) was used to avoid sharp, high-SF borders. The amplitude of *f* and *2f* was set to 66% and 33% contrast, respectively. To account for contrast reductions under AO-induced blur and maintain the *f*/*2f* contrast ratio (Exp.1,2), the amplitude of the two components was adjusted using corresponding MTF values. A fundamental frequency of 3 cycles/deg (*2f*: 6 cycles/deg) was used in all experiments (except Exp.S1).

### Phase perception task

On each trial, participants reported whether a compound grating stimulus presented foveally for 80ms appeared upwards or downwards (**Fig.3B,C**). No feedback was provided. All participants were first familiarized with the task using compound gratings presented until response. We computed the percent of upwards trials as a function of the *f*-*2f* relative phase. We used the Palamedes Toolbox (Prins and Kingdom, 2018) to fit data with a Cumulative Normal psychometric function using maximum likelihood estimation, and estimate the point-of-subjective equality (PSE)–i.e., the relative phase at which stimuli were perceived “aligned”. Non-parametric bootstrapping was used to compute 95%-CIs of PSE estimates and assess differences in PSE estimates. We randomly resampled individual trials with replacement to generate and fit resampled datasets. This procedure of resampling and refitting was repeated 10,000 times to generate bootstrap distributions of PSE estimates. Goodness of fit was verified using log likelihood with a global maximum successfully found for all fits.

### Visual acuity

Multiple visual acuity (VA) thresholds (10±6 on average) were measured for each participant under AO correction using a 4-AFC letter orientation task. On each trial, participants reported the orientation of a high-contrast, tumbling E acuity letter (Sloan) presented for 250ms. Letter size was adjusted from trial-to-trial using the QUEST staircase method (Watson and Pelli, 1983) to estimate 62.5%-correct thresholds (40 trials/staircase). Note that all included thresholds were obtained from successfully converged staircases, and were highly consistent within each observer.

### Optical theory predictions

The transfer function of the eye, like any optical system, can be characterized by the Modulation Transfer Function (MTF) and Phase Transfer Function (PTF), which specify the amplitude reduction and phase shift as a function of the signal SF, respectively. To obtain the MTF and PTF, we first compute the generalized pupil function containing the eye’s aberrations, as follows:

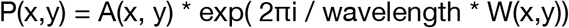

where P(x,y) is the pupil function at x,y Cartesian coordinates, A(x,y) the amplitude function, and W(x,y) the wavefront aberrations. This function is then used to calculate the point spread function (PSF) and optical transfer function (OTF) using 2-dimensional Fourier Transform (fft2):

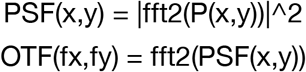

The OTF can further be decomposed into the MTF and PTF by taking the modulus (magnitude) and phase of the complex OTF:

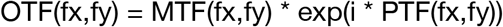

Predicted shifts in relative phase between *f* and *2f* components were computed using PTF values for various amounts of vertical coma (**Fig.4B**). These functions were used to generate simulated images for specific optical conditions, with or without compensating for the impact of MTF or PTF (**Fig.1**).

### Phase shift validation

Predicted phase shifts under specific amounts of vertical coma selected for testing in Exp.1 were validated directly from the AOVS (**Fig.4C**). A foldable mirror was inserted into the path to redirect it onto a CCD sensor and take measurements of the display. A collimated single mode diode laser was used to provide a signal to the Shack-Hartmann sensor and induce aberrations without the eye’s optics. Gratings of varying *f*-*2f* frequencies were displayed under AO-induced vertical coma ranging from -0.5 µm to +0.5 µm. Ten images were collected for each condition and averaged together. Segments of *f* and *2f* images were fit with cosine waves of variables amplitude, period, phase and vertical translation. The relative phase between the *f* and *2f* was then computed from these fits to validate predictions from our theoretical simulations.

### Blur-induced shifts in perceived phase (Exp.1)

Observers judged the appearance of compound gratings (*f*: 3 cycles/deg; relative phase: -90º to +90º, 10º steps) under nine conditions of AO-induced vertical coma (0, ±0.2, ±0.3, ±0.4, ±0.5µm). We selected this range of values to ensure that the magnitude of observed phase shifts could be effectively evaluated using the compound grating appearance task (i.e., within ±90º in relative phase, for reliable PSE estimation). As described above, the amplitudes of *f* and *2f* components were adjusted according to MTF values to account for blur-induced contrast reductions and maintain the *f*/*2f* contrast ratio. PSE estimates were compared to optical theory predictions (**Fig.4D-F**). Each condition was tested on separate blocks counterbalanced across sessions. In the MTF-adjusted control condition (**Fig.4E**), we tested the impact of blur-induced contrast reductions on perceived phase by testing participants under AO correction using compound stimuli whose contrast and relative phase were adjusted according to both MTF and PTF values for ±0.5µm vertical coma (i.e., phase offset: ±52.3º). 95%-CIs are provided to assess changes in PSEs across conditions. A total of 6097 trials was collected on average per observer (∼553 trials for each of the eleven tested conditions). Additional data was collected using higher-SF compound gratings (*f*: 4 cycles/deg) and positive vertical coma (**Fig.S2**).

### Short-term blur adaptation (Exp.2)

We assessed the effects of sustained exposure to vertical coma on perceived phase (**Fig.5**). Participants judged compound gratings (*f*: 3 cycles/deg) with relative phase ranging from -90º to +90º (15º steps). Each experimental session (**Fig.5A**) consisted of 1599 trials divided into 41 successive blocks (39 trials each) across 3 periods: baseline, adaptation, and post-adaptation. The baseline (pre-adaptation) consisted of 8 blocks (312 trials; ∼8 min) collected under AO correction (*PSE*_*predicted*_: 0º). During blur adaptation, observers were exposed to AO-induced vertical coma (−0.4µm; *PSE*_*predicted*_: +29.5º) over 21 blocks (819 trials; ∼70 min). The first 3 blocks (117 trials; ∼3 min) only contained compound stimuli to estimate the initial blur-induced shift in PSE. The following 18 adaptation blocks (702 trials; ∼67 min) contained natural images (e.g., animals, buildings, walls, humans) and checkerboard textures presented before each compound stimulus, serving as perceptual cues the visual system could adapt to in the presence of blur (**Fig.5B,C**). Each trial started with 6-12 adaptation stimuli, each presented for 240ms and separated by 13ms. After a 253-ms delay, a compound stimulus was presented for 80ms. Note that adaptation stimuli changed rapidly to homogenize local light adaptation and minimize afterimages, as done in previous studies(Elliott et al., 2011, 2007; Sawides et al., 2011a). Each adaptation stimulus had an equal probability to be a natural image or a checkerboard stimulus. Natural images were randomly selected from ∼700 natural grayscale images and could not repeat within the same trial. Checkerboard stimuli had random square size (3,4,5 or 6 square width) and phase. Immediately following blur adaptation, AO-correction was switched back on and 12 post-adaptation blocks (468 trials; ∼12min) were collected with only compound gratings presented, as in the baseline. Participants were allowed to take short breaks (∼1-2 min each) away from the bite-bar at specific moments within each session (**Fig.5A**). To minimize the impact of these interruptions on the adaptation speed or depth, participants were asked to stay in the experimental room under the same lighting conditions. During blur adaptation, the first trial following a break started with top-up adaptation (60 adaptation images)(**Fig.5B**). The control condition was identical to the blur adaptation condition, except that participants were always tested under full AO-correction. Instead of inducing blur, a phase offset of +29.5º was added to compound stimuli, mimicking the phase shift under -0.4µm of vertical coma. Thus, participants’ responses were similarly biased during the adaptation period. However, there was no clear phase disruption the visual system could detect and adapt to as adaptation stimuli were presented under full AO correction.

Each participant performed at least two experimental sessions on separate days. To better characterize changes in perceived phase across time, PSE estimates at a specific timepoint *t* (i.e., block number) were computed by binning trials from adjacent blocks (±1; i.e., trials from *t-1, t*, and *t+1*). This binning procedure was adjusted to ensure not to bin trials across different experimental periods (baseline, adaptation and post-adaptation), for which trials from only two blocks were binned together. Importantly, our results did not depend on this binning procedure (**Fig.S3**). 95%-CIs were provided to assess changes in PSEs across time. In order to better visualize PSE changes relative to baseline, the mean of PSE estimates during baseline was subtracted from all PSE estimates, correcting for small individual biases and treating 0º as reference.

### Impact of long-term exposure to poor optical quality (Exp.3)

We assessed the effects of long-term (i.e., months to years) exposure to poor optical quality on phase perception. We measured perceived phase in patients with keratoconus (KC)–a corneal disease causing severe optical blur in neurotypically-developed adults (**Fig.6A**). KC results in large amounts of HOAs, particularly of vertical coma that primarily affects resolution along the vertical axis (i.e., horizontally-oriented signals). As previously detailed (Barbot et al., 2021; Ng et al., 2022; Sabesan et al., 2017; Sabesan and Yoon, 2010, 2009; Zheleznyak et al., 2016), our AOVS is able to correct all optical aberrations and estimate visual functions under similar, aberration-free conditions in both healthy and KC eyes. We measured PSE estimates under full AO correction in eleven control participants with healthy eyes and six KC patients (**Fig.6B,S5**). Relative to the optical quality of control participants, habitual optical quality in KC eyes was severely degraded despite their everyday optical correction (tRMS: 1.06±0.31µm in controls, 4.04±2.3µm in KCs), particularly when looking at the amount of uncorrected HOAs (hRMS: 0.38±0.13µm in controls, 2.47±1.33µm in KCs). Poorer habitual optical quality was associated with stronger alterations of visual functions (**Fig.6,S6**). Note that all participants were tested across ±90º range in relative phase, but the exact values used slightly varied mainly due to time constraints, with either 19 phase levels (10º equal steps), 13 phase levels (15º equal steps), or 9 phase levels (±90º,±45º,±15º,±7.5º, and 0º). A total of ∼900 trials was collected on average per participant.

## Article information

### Author contributions

AB and GY conceived and designed the study. AB, JTP and CLN performed the experiment. AB, JTP and GY analyzed the data. AB and GY wrote the paper, and all authors commented on it.

### Data availability

Data will be publicly available from the OSF database (https://osf.io/ug8mr/).

## Acknowledgment

This work was supported by NIH/NEI grants RO1-EY014999, P30-EY001319 (core grant to Center for Visual Science, University of Rochester), Research to Prevent Blindness, and Schmitt Program on Integrative Neuroscience (University of Rochester). We thank Dr. Tara Vaz, OD and Olga Pikul for helping with participant screening and recruitment. We also thank Duje Tadin, Randolph Blake and Martin Banks for comments on an earlier version of the manuscript.

## References

Amano S, Amano Y, Yamagami S, Miyai T, Miyata K, Samejima T, Oshika T. 2004. Age-related changes in corneal and ocular higher-order wavefront aberrations. Am J Ophthalmol 137. doi:10.1016/j.ajo.2004.01.005

Anzai A, Ohzawa I, Freeman RD. 1997. Neural mechanisms underlying binocular fusion and stereopsis: position vs. phase. Proc Natl Acad Sci U S A 94:5438–5443. doi:10.1073/pnas.94.10.5438

Aronov D, Reich DS, Mechler F, Victor JD. 2003. Neural coding of spatial phase in V1 of the macaque monkey. J Neurophysiol 89:3304–3327. doi:10.1152/jn.00826.2002

Artal P, Chen L, Fernandez EJ, Singer B, Manzanera S, Williams DR. 2004. Neural compensation for the eye’s optical aberrations. J Vis 4:281–287. doi:10:1167/4.4.44/4/4 [pii]

Atkinson J, Campbell FW. 1974. The effect of phase on the perception of compound gratings. Vision Res 14:159–162. doi:10.1016/0042-6989(74)90096-0

Badcock DR. 1984. How do we discriminate relative spatial phase? Vision Res 24:1847–1857. doi:10.1016/0042-6989(84)90017-8

Bao M, Engel SA. 2012. Distinct mechanism for long-term contrast adaptation. PNAS 109:5898–5903. doi:10.1073/pnas.1113503109

Barbot A, Park WJ, Ng CJ, Zhang RY, Huxlin KR, Tadin D, Yoon G. 2021. Functional reallocation of sensory processing resources caused by long-term neural adaptation to altered optics. Elife 10. doi:10.7554/eLife.58734

Bennett PJ, Banks MS. 1987. Sensitivity loss in odd-symmetric mechanisms and phase anomalies in peripheral vision. Nature 326:873–876. doi:10.1038/326873a0

Bex PJ, Solomon SG, Dakin SC. 2009. Contrast sensitivity in natural scenes depends on edge as well as spatial frequency structure. J Vis 9:1–1. doi:10.1167/9.10.1

Braddick OJ, Atkinson J, Wattam-Bell JR. 1986. Development of the discrimination of spatial phase in infancy. Vision Res 26. doi:10.1016/0042-6989(86)90103-3

Burr DC. 1980. Sensitivity to spatial phase. Vision Res 20:391–396.

Elliott SL, Georgeson MA, Webster MA. 2011. Response normalization and blur adaptation: Data and multiscale model. J Vis 11:7–7. doi:10.1167/11.2.7

Elliott SL, Hardy JL, Webster MA, Werner JS. 2007. Aging and blur adaptation. J Vis 7:8–8. doi:10.1167/7.6.8

Freeman J, Simoncelli EP. 2011. Metamers of the ventral stream. Nat Neurosci 14:1195–1201. doi:nn.2889 [pii] 10.1038/nn.2889

Georgeson MA, Sullivan GD. 1975. Contrast constancy: deblurring in human vision by spatial frequency channels. J Physiol 252:627–656. doi:10.1113/jphysiol.1975.sp011162

Hastings GD, Schill AW, Hu C, Coates DR, Applegate RA, Marsack JD. 2021. Orientation-specific longterm neural adaptation of the visual system in keratoconus. Vision Res 178. doi:10.1016/j.visres.2020.10.002

Henriksson L, Hyvarinen A, Vanni S. 2009. Representation of cross-frequency spatial phase relationships in human visual cortex. J Neurosci 29:14342–14351. doi:10.1523/JNEUROSCI.3136-09.2009

Hyvarinen A, Gutmann M, Hoyer PO. 2005. Statistical model of natural stimuli predicts edge-like pooling of spatial frequency channels in V2. BMC Neurosci 6:12. doi:10.1186/1471-2202-6-12

Kompaniez EJ, Sawides L, Marcos S, Webster MA. 2013. Adaptation to interocular differences in blur. J Vis 13:1–14. doi:10.1167/13.6.19

Kovesi P. 2000. Phase congruency: a low-level image invariant. Psychol Res 64:136–148.

Liang J, Williams DR, Miller DT. 1997. Supernormal vision and high-resolution retinal imaging through adaptive optics. J Opt Soc Am A Opt Image Sci Vis 14:2884–2892.

McCormick GJ, Porter J, Cox IG, MacRae S. 2005. Higher-order aberrations in eyes with irregular corneas after laser refractive surgery. Ophthalmology 112. doi:10.1016/j.ophtha.2005.04.022

Mechler F, Reich DS, Victor JD. 2002. Detection and discrimination of relative spatial phase by V1 neurons. J Neurosci 22:6129–6157. doi:20026297

Mon-Williams M, Tresilian JR, Strang NC, Kochhar P, Wann JP. 1998. Improving vision: neural compensation for optical defocus. Proc Biol Sci 265:71–77. doi:10.1098/rspb.1998.0266

Morrone MC, Burr DC. 1988. Feature detection in human vision: a phase-dependent energy model. Proc R Soc Lond B Biol Sci 235:221–245. doi:10.1098/rspb.1988.0073

Morrone MC, Burr DC, Spinelli D. 1989. Discrimination of spatial phase in central and peripheral vision. Vision Res 29:433–445.

Murray S, Bex PJ. 2010. Perceived Blur in Naturally Contoured Images Depends on Phase. Front Psychol 1. doi:10.3389/FPSYG.2010.00185

Ng CJ, Blake R, Banks MS, Tadin D, Yoon G. 2021. Optics and neural adaptation jointly limit human stereovision. Proceedings of the National Academy of Sciences 118:e2100126118. doi:10.1073/pnas.2100126118

Ng CJ, Sabesan R, Barbot A, Banks MS, Yoon G. 2022. Suprathreshold Contrast Perception Is Altered by Long-term Adaptation to Habitual Optical Blur. Invest Ophthalmol Vis Sci 63:6. doi:10.1167/iovs.63.11.6

Oppenheim A, Lim J. 1981. The importance of phase in signals. Proc IEEE 69:529–541.

Pantanelli S, MacRae S, Jeong TM, Yoon G. 2007. Characterizing the wave aberration in eyes with keratoconus or penetrating keratoplasty using a high-dynamic range wavefront sensor. Ophthalmology 114:2013–2021. doi:S0161-6420(07)00072-3 [pii] 10.1016/j.ophtha.2007.01.008

Perna A, Tosetti M, Montanaro D, Morrone MC. 2008. BOLD response to spatial phase congruency in human brain. J Vis 8:15 1–15. doi:10.1167/8.10.15

Prins N, Kingdom FAA. 2018. Applying the Model-Comparison Approach to Test Specific Research Hypotheses in Psychophysical Research Using the Palamedes Toolbox. Front Psychol 9:1250. doi:10.3389/fpsyg.2018.01250

Radhakrishnan A, Dorronsoro C, Sawides L, Webster MA, Marcos S. 2015. A cyclopean neural mechanism compensating for optical differences between the eyes. Current Biology. doi:10.1016/j.cub.2015.01.027

Rajeev N, Metha A. 2010. Enhanced contrast sensitivity confirms active compensation in blur adaptation. Invest Ophthalmol Vis Sci 51:1242–1246. doi:10.1167/iovs.09-3965

Roorda A. 2011. Adaptive optics for studying visual function: a comprehensive review. J Vis 11. doi:10.1167/11.5.6

Rosenfield M, Hong SE, George S. 2004. Blur adaptation in myopes. Optom Vis Sci 81:657–662.

Rouger H, Benard Y, Gatinel D, Legras R. 2010. Visual Tasks Dependence of the Neural Compensation for the Keratoconic Eye’s Optical Aberrations. J Optom 3:60–65.

Sabesan R, Ahmad K, Yoon G. 2007. Correcting highly aberrated eyes using large-stroke adaptive optics. Journal of Refractive Surgery 23:947–952.

Sabesan R, Barbot A, Yoon G. 2017. Enhanced neural function in highly aberrated eyes following perceptual learning with adaptive optics. Vision Res 132:78–84. doi:10.1016/j.visres.2016.07.011

Sabesan R, Yoon G. 2010. Neural compensation for long-term asymmetric optical blur to improve visual performance in keratoconic eyes. Invest Ophthalmol Vis Sci 51:3835–3839. doi:iovs.09-4558 [pii] 10.1167/iovs.09-4558

Sabesan R, Yoon G. 2009. Visual performance after correcting higher order aberrations in keratoconic eyes. J Vis 9:6 1–10. doi:10.1167/9.5.69/5/6 [pii]

Sawides L, de Gracia P, Dorronsoro C, Webster M, Marcos S. 2011a. Adapting to blur produced by ocular high-order aberrations. J Vis 11. doi:11.7.21 [pii] 10.1167/11.7.21

Sawides L, de Gracia P, Dorronsoro C, Webster MA, Marcos S. 2011b. Vision is adapted to the natural level of blur present in the retinal image. PLoS One 6:e27031. doi:10.1371/journal.pone.0027031

Sawides L, Marcos S, Ravikumar S, Thibos L, Bradley A, Webster M. 2010. Adaptation to astigmatic blur. J Vis 10:22. doi:10.1167/10.12.22

Venkataraman AP, Winter S, Unsbo P, Lundström L. 2015. Blur adaptation: Contrast sensitivity changes and stimulus extent. Vision Res 110:100–106. doi:10.1016/J.VISRES.2015.03.009

Vinas M, Sawides L, de Gracia P, Marcos S. 2012. Perceptual Adaptation to the Correction of Natural Astigmatism. PLoS One 7:e46361. doi:10.1371/JOURNAL.PONE.0046361

Wang Z, Simoncelli EP. 2004. Local Phase Coherence and the Perception of Blur. Advances in Neural Information Processing Systems 16.

Watson AB, Pelli DG. 1983. QUEST: a Bayesian adaptive psychometric method. Percept Psychophys 33:113–120.

Webster MA. 2017. Blur Adaptation and Induction. The Oxford Compendium of Visual Illusions 756–760. doi:10.1093/ACPROF:OSO/9780199794607.003.0110

Webster MA. 2015. Visual Adaptation. Annu Rev Vis Sci 1:547–567. doi:10.1146/annurev-vision-082114-035509

Webster MA, Georgeson MA, Webster SM. 2002. Neural adjustments to image blur. Nat Neurosci 5:839– 840. doi:10.1038/nn906nn906 [pii]

Wichmann FA, Braun DI, Gegenfurtner KR. 2006. Phase noise and the classification of natural images. Vision Res 46:1520–1529. doi:10.1016/J.VISRES.2005.11.008

Williams PE, Shapley RM. 2007. A dynamic nonlinearity and spatial phase specificity in macaque V1 neurons. J Neurosci 27:5706–5718. doi:10.1523/JNEUROSCI.4743-06.2007

Yoon G, Jeong TM, Cox IG, Williams DR. 2004. Vision improvement by correcting higher-order aberrations with phase plates in normal eyes. J Refract Surg 20:S523–7.

Yoon GY, Williams DR. 2002. Visual performance after correcting the monochromatic and chromatic aberrations of the eye. J Opt Soc Am A Opt Image Sci Vis 19:266–275.

Zhang F, Jiang W, Autrusseau F, Lin W. 2014. Exploring V1 by modeling the perceptual quality of images. J Vis 14:26–26. doi:10.1167/14.1.26

Zheleznyak L, Barbot A, Ghosh A, Yoon G. 2016. Optical and neural anisotropy in peripheral vision. J Vis 16:1. doi:10.1167/16.5.1

